# Membrane tension can enhance adaptation to maintain polarity of migrating cells

**DOI:** 10.1101/2020.04.01.020289

**Authors:** C. Zmurchok, J. Collette, V. Rajagopal, W. R. Holmes

**Affiliations:** Vanderbilt University; University of Melbourne

**Keywords:** adaptation, mechanosensing, Rac GTPase, cell polarity, neutrophil, tension

## Abstract

Migratory cells are known to adapt to environments that contain wide-ranging levels of chemoattractant. While biochemical models of adaptation have been previously proposed, here we discuss a different mechanism based on mechanosensing, where the interaction between biochemical signaling and cell tension facilitates adaptation. We describe and analyze a model of mechanochemical-based adaptation coupling a mechanics-based physical model of cell tension coupled with the wave-pinning reaction-diffusion model for Rac activity. Mathematical analysis of this model, simulations of a simplified 1D cell geometry, and 2D finite element simulations of deforming cells reveal that as a cell protrudes under the influence of high stimulation levels, tension mediated inhibition of GTPase signaling causes the cell to polarize even when initially over-stimulated. Specifically, tension mediated inhibition of GTPase activation, which has been experimentally observed in recent years, facilitates this adaptation by countering the high levels of environmental stimulation. These results demonstrate how tension related mechanosensing may provide an alternative (and potentially complementary) mechanism for cell adaptation.

**Statement of Significance:** Migratory cells such as human neutrophils encounter environments that contain wide-ranging levels of chemoattractant. In order to move, these cells must maintain an organized front-rear signaling polarity despite this wide variation in environmental stimuli. Past research has demonstrated a number of biochemical based mechanisms by which cells adapt to variable signal levels. Here we demonstrate that the interplay between Rho GTPase signaling and tension mediated feedbacks *may* provide an alternative mechanochemical mechanism for adaptation to high levels of signaling.

## Introduction

Human neutrophils and other migratory cells are able adapt to environments that contain wide-ranging levels of chemoattractant to maintain polarity. How does this adaptation occur? Models of adaptation based on biochemical processes such as receptor occupancy or receptor arrangement [1] have long been proposed. Here we demonstrate that well-established mechanochemical interactions between Rac signaling and protrusion related changes in cell tension can facilitate adaptation to maintain polarity when cells are exposed to wide-ranging levels of chemoattractant.

In the context of cell signaling, adaptation refers to the ability of cells to adjust to high levels of signaling or stimulation. Most signal response systems have a range of inputs over which they are sensitive to differences. When stimulus levels are sufficiently high to saturate the underlying regulatory system, the system loses its ability to distinguish signaling levels. For migration, cells must be able to adapt to higher stimulus levels as they proceed closer to the attractant source. In neutrophil migration, the Rho-family GTPase Rac rapidly polarizes to orient the cell in the direction of the stimulus [2, 3]. Interestingly, when these cells are subjected to high levels of stimulation, Rac first becomes highly activated throughout the cell and subsequently polarizes [3]. Thus, in response to high levels of stimulation, these cells first adapt and then subsequently polarize. It has been previously demonstrated that feedback from cell tension to GTPase activation causes competition between multiple cellular protrusions [4, 5] (Figure 1B), ensuring that the cell protrudes in only one direction at a time. Here we demonstrate that this feedback could also provide the cell with a form of mechanochemical adaptation to high, saturating levels of stimulation.

**Figure 1:**
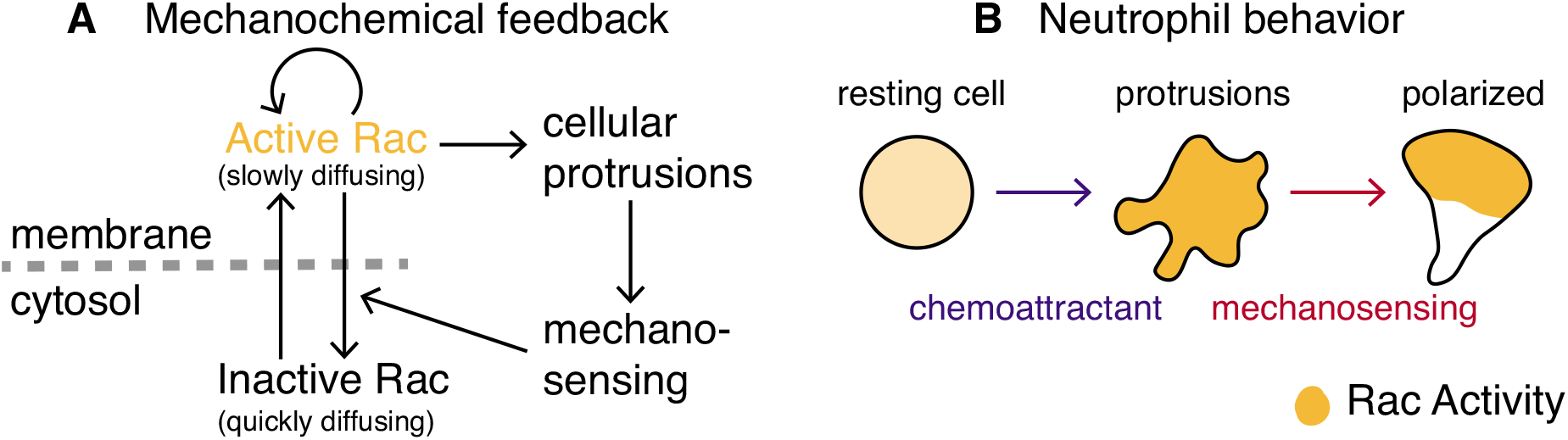
Overview of the mechanochemical feedback model and observed neutrophil behavior. **A** Schematic of mechanochemical feedback. Active (respectively inactive) Rac are assumed to be bound (respectively unbound) to the membrane and therefore slow (respectively fast) diffusing. Active Rac is autoactivated and causes cellular protrusions by activating downstream effectors. Cellular protrusions increase mechanosensing, which increases the deactivation rate of Rac. **B** Cartoon of observed neutrophil behavior in a chemoattractive environment. A resting neutrophil is washed in chemoattractant, causing multiple sites of protrusive activity. Subsequently the cell adapts to the high signaling environment and polarizes to form a single leading front. Panel B is adapted from Figure 3B of [5].

Our model mechanisms and results incorporate ideas related to adaptation, polarity, and tension regulation, each of which is well studied in its own right. Adaptation has been primarily studied from a biochemical perspective. Models traditionally devised to explain the run-and-tumble behavior of bacterial chemotaxis [6] typically incorporate either negative feedback loops [7] or incoherent feed forward loops [8] to achieve perfect adaptation to step changes in stimulus magnitude [9]. These ideas were later combined with Meinhardt’s concept of lateral inhibition for biological pattern formation [10] to explain how cells could adapt to environments and polarize in response to chemoattractant if the stimulus simultaneously triggered both a local excitatory activator and global inhibitor [11, 12]. This mechanism, known as LEGI (local excitation global inhibition), and variants have been extensively studied experimentally and theoretically [13–19]. However, not all forms of adaptation are biochemical in nature. Cells can adapt to sustained mechanical stresses [20], for example, and large-scale cytoskeletal structures are known to lead to adaptive responses in cell migration to stiffness gradients in the environment [21].

Similarly, polarity regulation itself has seen sustained interest for multiple decades [22]. Much (though not all) of this study has been dedicated to understanding the role of Rho GTPases, which regulate cytoskeletal remodeling, in polarity generation [23]. The wave-pinning model of polarity [24–26], which we will build on here, demonstrated that the conservative activation and inactivation dynamics of these proteins are sufficient to explain the broad characteristics of the polarization process. The cell polarity and cell adaptation literature have been bridged by coupled models that exhibit both adaptive and polarity characteristics [19]; however, such models are still biochemical in nature. More recently, models that couple single-cell mechanics and cytoskeletal mechanobiology with biochemistry have begun to explore the role of mechanochemical feedbacks on cell dynamics [3, 4, 27, 28]. However, the possible impact of cytoskeletal mechanics on adaptation during migration has yet to be explored.

Here, we build on the well-studied wave-pinning model of Rho GTPase dynamics [24–27, 29–33] to analyze the consequences of feedback between signaling, cell protrusion (which affects tension), and tension. We include three critical assumptions into this model. First, that Rac signaling (which is modeled using wave-pinning dynamics) promotes protrusion of the cell membrane through downstream activation of actin polymerization [23]. Second, protrusion dynamics related changes in cell surface area increase or decrease membrane tension. Third, membrane tension inhibits GTPase signaling [4, 5, 34–41]. We combine these assumptions into a moving-boundary mechanochemical model (Figure 1A) that incorporates reaction-diffusion partial differential equations (PDEs) for Rac activity in a moving domain and a continuum-based description of cellular mechanics that affect the domain.

To understand the resulting model, we first use Local Perturbation Analysis (LPA) [42, 43] and numerical bifurcation analysis to analyze how tension changes resulting from changes in cell size influence GTPase signaling. Results indicate that increases in tension cause the cell to transit from an overstimulated to a polarizable regime of the GTPase signaling model. To test whether this adaptive response induces polarization, we use 1D and 2D minimal models to describe cellular physics, and simulate the reaction-diffusion Rac signaling model in fully-moving cells. Our purpose here is not to develop a high fidelity model of cellular biophysics (see [44–55] for such models). Instead, our purpose is to use these models to assess how cell protrusions, tension, and GTPase signaling interact. Both 1D and 2D simulations demonstrate that the feedback between signaling and tension does lead to adaptation where the cell initially protrudes everywhere in response to a high stimulus but subsequently polarizes, as predicted. These results confirm that the feedback for establishment of polarity via membrane tension can be described using a wave-pinning model of Rac signaling, and more importantly that mechanosensing may (in addition to biochemical forms of adaptation) facilitate adaptation of migratory cells to high signaling levels.

## Models and methods

We will develop and analyze a model of the feedbacks between Rac signaling, cell size change, and tension changes. The purpose here is to use this model to study how these feedbacks may influence a cell’s response to large stimuli. More specifically, we use this model to explore the hypothesis that these feedbacks provide a mechanochemical form of adaptation.

Here we briefly outline the core elements of the model and analysis approach. We provide further details in subsequent sections. The model (illustrated in Figure 1A) builds on the well-studied wave-pinning model of Rho GTPase dynamics [24–27, 29–33] to integrate tension effects resulting from changes in cell surface area. At a broad level, this model incorporates the cycling of Rac between activated (membrane bound) and inactivated (cytosolic) states, Rac-mediated tension production (via cell protrusion), and tension-mediated inhibition of Rac activation (see below for further detail), all of which are well-established in neutrophil cells [4, 5, 34–41].

Since addressing the questions of interest here involves non-linear interactions between signaling, shape change, and mechanics, we take several steps to analyze the dynamics of this system, with each building in additional complexity. First, we use the Local Perturbation Analysis (a form of spatio-temporal bifurcation analysis) to approximately map out the parametric dynamics of the system. Second, we study the effects of tension and cell size changes over time using 1D fixed domain simulations of the spatial PDE model where tension and length are treated as parameters that we manually vary over time. This allows us to manually simulate tension effects to understand their potential effects on signaling dynamics. Third, we simulate a 1D moving-boundary version of the model where Rac promotes protrusion, leading to cell size changes and tension effects. Finally, we simulate a 2D moving-boundary version of this model coupling continuum mechanics with Rac signaling.

### Reaction-diffusion Rac GTPase signaling model

We first describe the core model of Rac signaling that will be responsible for regulating protrusion in this model. We adopt the well-studied wave-pinning model [24–27, 29–33] of GTPase activity. We consider Rac activity in a 1D or 2D domain with no-flux boundary conditions at the cell periphery and model the activity of both membrane-bound active Rac (*R*(*x, t*)) and freely-diffusing cytosolic inactive Rac (*R_i_*(*x, t*)) forms as illustrated in the left part of Figure 1A. The non-dimensionalized PDEs describing these dynamics in a stationary domain are:

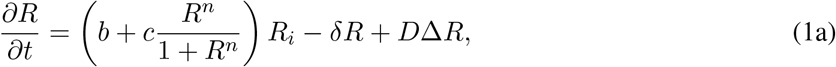

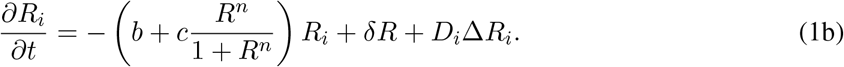

The term in parentheses is a standard Hill function representation of auto-activation kinetics with *b* representing a basal activation rate and *c* the magnitude of auto-activation (i.e. positive feedback). The parameter *δ* is the deactivation rate, and *D* and *D_i_* are the diffusion coefficients of the active and inactive forms, respectively. The ratio *D/D_i_* is large because membrane-bound diffusion is slower than cytoplasmic diffusion. Since Rac cycles between inactive and active forms with no-flux boundary conditions, the total amount is conserved as a function of time: *∫ R*(*x, t*) + *R_i_*(*x, t*) *dx = R_T_*. Full model and non-dimensionalization details are in the Supporting Material.

### External chemoattractant stimulation and feedback from tension

The two critical parameters of this model that we will focus on are the basal activation rate *b* and the inactivation rate *δ*. The basal activation rate will be used to encode the level of external stimulation applied to the cell. For example, application of a uniform stimulus will be modelled as an increase in the parameter *b* as the Rac signaling pathway becomes activated upon external stimulation. The *δ* parameter will be used to describe tension (*T*) mediated inhibition of signaling:

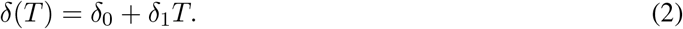

In cases where we are not explicitly modeling tension, we will describe the effects of tension through changes in the parameter *δ*. For example, an increase in tension will be assumed to increase *δ*. Other functional forms (e.g., Hill functions) of *δ*(*T*) could of course be used. We use a linear function here for simplicity and to reduce the number of extraneous parameters in the model.

### Local Perturbation Analysis, bifurcation analysis, and steady-state behavior

To approximately determine how stimulus level (*b*) and tension strength (*δ*) jointly affect the spatial dynamics of this system, we used Local Perturbation Analysis (LPA) to analyze the behavior of the PDE model in the *bδ*-plane. The LPA is an approximate asymptotic analysis that predicts whether the model (Equation (1)) will respond to a spatial perturbation (either small or large) and give rise to a spatial pattern formation. More specifically, it helps identify regions of parameter space associated with both linear instabilities (i.e., Turing regions) and stimulus induced patterning (where large stimuli are required for a response). The LPA has been demonstrated to be an effective analysis tool for studying GTPase and other systems [29, 31, 33, 56–60].

In practice, this method reduces the PDE system (Equation (1)) to an ordinary differential equation (ODE) system that approximates the dynamics of an asymptotically localized perturbation. By mathematically analyzing how the model system responds to this type of mathematically convenient perturbation, we can efficiently assess whether the model will give rise to spatial pattern formation and map its parameter space. This is carried out by studying the reduced LPA ODE system with numerical bifurcation software (Matcont [61]) to determine the perturbed system’s behavior in different parameter regimes (Figure 2A). Full details of the bifurcation analysis of the LPA ODE system can be found in the Supporting Material.

**Figure 2:**
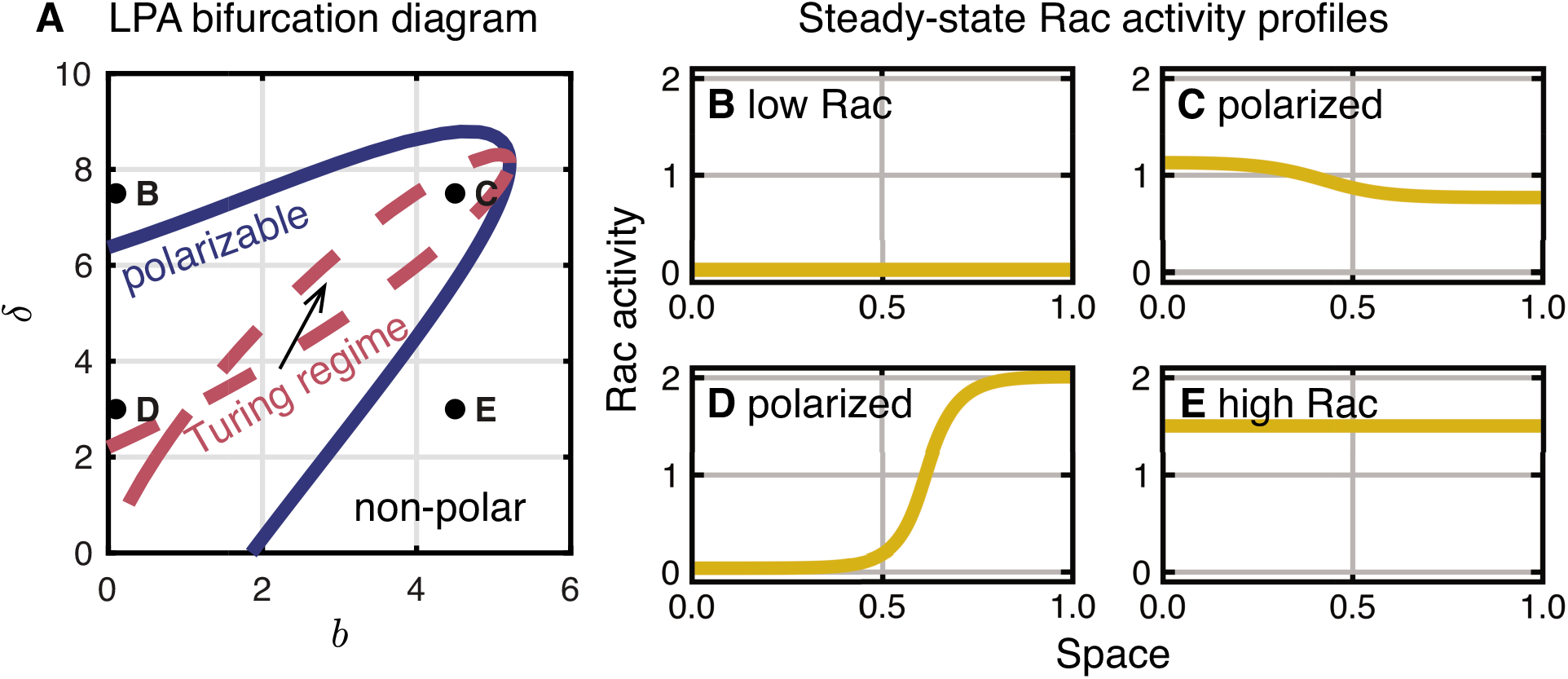
Parameter regimes of polar and non-polar behavior and steady-state PDE solutions on a stationary domain. **A** Local perturbation analysis (LPA) reveals regimes of non-polar, stimulus-induced polarizable, and Turing unstable steady-state (i.e., polarizable) cell behavior in the *bδ*-plane. Dots correspond to parameters for steady-state Rac activity profiles (solid curves) in B-E. **B-E** Steady-state simulated solutions to the PDEs. Initial conditions *R*(*x*, 0) = 0 except *R*(*x*, 0) = 2 for *x* > 0.9 are used in B, D, and E. For C, *R*(*x*, 0) = 1 + sin(4*πx*)/10 is used. Spatially homogeneous initial conditions were used for the inactive amount to ensure the total Rac (*R_T_*) is preserved: 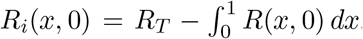. **B** Low Rac activity *b* = 0.1, *δ* = 7.5. **C** Polarized Rac activity *b* = 0.1, *δ* = 3. **D** Polarized Rac activity in the Turing regime *b* = 4.5, *δ* = 7.5. **E** High Rac activity *b* = 4.5, *δ* = 3. Other parameters are *D* = 0.01, *D_i_* = 10, *γ* = 5, *n* = 6, and *R_T_* = 2.

### 1D mechanochemical model

Before moving to a full two-dimensional changing domain model of the cell, we first develop a simpler onedimensional changing domain model to provide a preliminary analysis of the effects of tension and signaling feedbacks. To interrogate the mechanosensing-adaptation hypothesis in a moving, 1D cell, we couple the wave-pinning model for Rac activity to a simple spring-like mechanical model of the deforming cell.

This minimal 1D deforming cell model consists of two components: (1) an over-damped, elastic spring model of the cell and (2) an adapted version of the Rac model to appropriately account for the changing domain. These are coupled with two conditions. First, we assume that active Rac at the cell ends can exert a protrusive force on the cell that opposes the restoring linear elastic spring force. This protrusive force depends sigmoidally on Rac activity at the endpoints. Second, the deactivation rate *δ* increases linearly with tension according to Equation (2). Here, we define tension to be the difference between the cell length *L*(*t*) and rest-length ℓ_0_: *T = L*(*t*) − ℓ_0_. Together, these assumptions result in a moving-boundary problem where the Rac activity is transported within a time-dependent domain Ω(*t*) = [*x*_−_(*t*), *x*_+_(*t*)]:

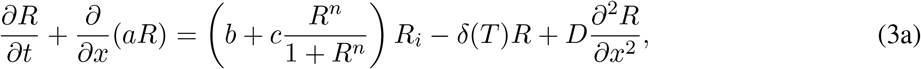

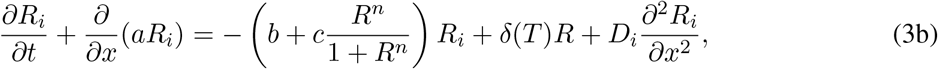

with no-flux boundary conditions. We can determine the velocity field *a*(*x, t*) throughout the cell body from the forces imposed at the cell ends since we assume that the cell changes length isotropically (as in [62–65]). This assumption means that as the cell length, *L*(*t*) = *x*_+_(*t*) − *x*_−_(*t*), changes, each Lagrangian volume element will change proportionally to the total length change (this is a simplifying approximation that will be relaxed in the 2D model to follow). This relation specifies the flow *a* so that Rac is transported in the domain accordingly. Full model details can be found in the Supporting Material.

We assume that the movement of the cell’s endpoints depends on a linear elastic restoring force and protrusive forces determined by Rac activity at the endpoints:

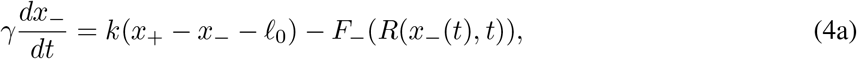

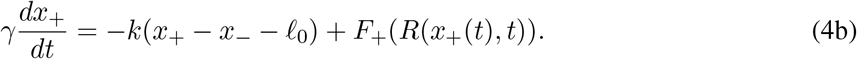

Here, *γ* is the viscosity, *k* is the spring constant, ℓ_0_ is the rest-length of the spring and the functions *F*_±_ describe the protrusive forces oriented outward from the cell from *R*. Note that inertial effects are ignored as appropriate for modeling cell motion. We use a smoothed Heaviside function for the Rac-dependence force function *F*_±_(*R*):

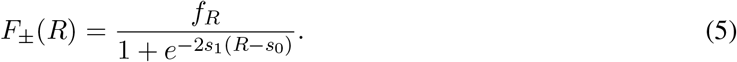

The parameters *s*_1_ and *s*_0_ control the sharpness and location of the switch, respectively, and *f_R_* > 0 is the magnitude of the force due to Rac activity.

### 2D mechanochemical model

We extended the 1D mechanochemical model study to a 2D domain by simulating coupled mechanics and reaction diffusion equations on a 2D circular deformable finite element mesh geometry. As a consequence, the domain can freely deform in response to Rac activity and mechanics (domain changes are no longer isotropic). We assume that the diffusion of active and inactive Rac is isotropic in the *x* and *y* directions and leave the reactions unchanged. The 2D deforming continuum mechanics model is described by the first order dynamic stress equilibrium equations and the linear viscoelastic stress-strain relationship. The kinematic relationships between the undeformed and deformed configurations are described by small, infinitesimal strains. These relationships are defined in terms of the stress ***σ***, strain ***ε***, and displacement ***u*** fields as:

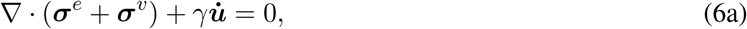

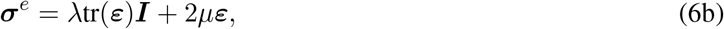

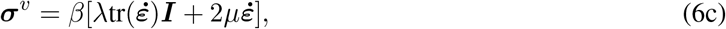

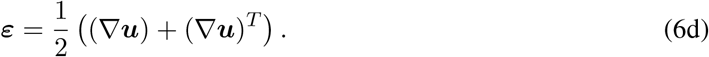

In the dynamic stress equilibrium equation (Equation (6)), *γ* is the Stokes damping coefficient and represents the viscous forces in the extracellular environment that resist cell motion. The contribution of intracellular mechanical forces to stress equilibrium are captured by Kelvin-Voigt viscoelastic stresses ***σ*** = ***σ**^e^* + ***σ**^υ^* (Equation (6a)). We assume that the elastic stress ***σ**^e^* is linear defined by the Lamé Pamameters λ, *μ* (Hooke’s law; Equation (6b)), and the viscous stress ***σ**^υ^* follows an isotropic linear constitutive model defined by a viscous parameter *β*, which is related to the first and second coefficients of viscosity *β*_λ_, *β_μ_* (Equation (6c)). We use the infinitesimal, small-strain approximation to describe the kinematic relationships between the undeformed and deformed configurations of the 2D domain (Equation (6d)).

The mechanical model is coupled to Rac activity through the same conditions as in the 1D context but with three modifications. First, we assume the protrusive force from active Rac acts normal to the membrane boundary at any given point. Second, we assume that the deactivation rate now depends on a proxy for 2D tension, *T* = *A*(*t*) − *A*_0_, the difference between the area *A*(*t*) and rest-area *A*_0_. Finally, we account for the deforming mechanics domain in the solution of the reaction diffusion system as follows.

The kinematic relationship between an initial and deformed geometry in the mechanics model is described by the deformation gradient tensor 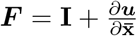 (where **I** is the identity tensor) and its determinant *J* = det(***F***). This relationship can be used to map the spatial and temporal differential operators, and allows the reaction-diffusion equations to be solved on a stationary reference domain with fixed coordinates 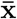:

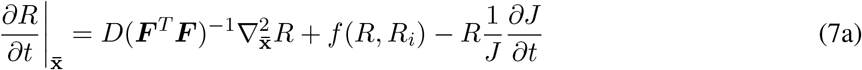

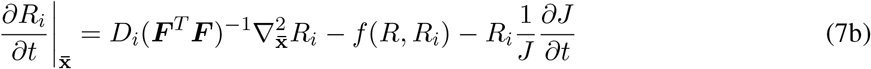

Full model details can be found in the Supporting Material.

### Numerical methods

We used a method of lines discretization and the Python function odeint from SciPy [66] to integrate the ODE system resulting from the discretization of the 1D PDE model described in Equation (2) and Matplotlib [67] for visualization. We generated the steady-state Rac activity profiles in Figure 2(b) with this method using *N* = 1000 spatial grid points and integrated until time *t* = 100. We used the same method for all 1D PDE numerics; however, we adapted the scheme for time-dependent parameters and to simulate the mechanochemical model. To do so, we first transformed the moving-boundary problem to a stationary-domain. Next, we solved the stationary-domain problem while simultaneously solving additional ODEs for the location of the cell’s endpoints. By reversing the transformation we can obtain the full moving-boundary solution. This method is similar to that used in other investigation of pattern formation on time-dependent domains [62–65].

We developed the finite element implementation of the 2D model using a customized reaction-diffusion module and linear-viscoelasticity module within an open source finite element modeling package, Open-CMISS [68]. The reaction-diffusion and mechanics equations were solved using a dynamic first-order backward Euler solver. The initial 2D domain for the finite element simulations is a circle of area 1 (radius 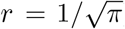). We used a quadratic simplex interpolation scheme to generate all meshes and adaptive refinement to increase number of elements closer to the boundary of the circular domain. This ensured that the finite element method adequately captured the reaction-diffusion dynamics and the high strain and stress gradients near the boundary. The resultant mesh contains 2, 358 elements and 4, 837 degrees of freedom for 2 chemical species in the reaction-diffusion system and for displacements in the mechanics equations.

We validated the numerical methods in both 1D and 2D by checking for Rac mass conservation over time. We assessed whether a simulated cell was polarized by considering the difference between maximum and minimum Rac activity throughout the domain. If this difference is sufficiently high (> 0.1 in 1D and > 0.15 in 2D) we consider the cell polarized. See Supporting Material for full details.

### Parameters

We non-dimensionalized the Rac signaling PDEs (see Supporting Material for details). The Rac signaling parameters (*R_T_* = 2, *n* = 6 and *c* = 5, respectively) were chosen to be consistent with prior studies [31] of the wave-pinning motif that ensure a large polarizable region in the *bδ*-plane (basal-activation rate *b*; deactivation rate *δ*). In the 2D model, a small chemical gradient (*b*_1_ = 0.1) is applied to mimic a chemoattractant gradient and align polarization in the positive *x* direction. The large ratio of diffusion coefficients (*D/D_i_*) is based on known GTPase biology (slowly diffusing membrane-bound active forms *D* = 0.01 vs. quickly diffusing cytosolic inactive forms *D_i_* = 10 for 1D and *D_i_* = 100 for 2D). Using LPA, we mapped the behavior of the Rac signaling PDEs in the *bδ*-plane to determine the magnitude of the mechanochemical feedback parameters from tension (*δ*_0_ = 3 and *δ*_1_ = 1).

For the 1D mechanical model, we chose damping coefficient *γ* = 1, spring constant = 0.001, force magnitude from Rac *f_R_* = 0.001, and switch parameters, *s*_0_ = 1 and *s*_1_ = 10, so that changes in the cell length are smaller and slower compared to Rac activity. Additionally, these parameters were chosen so that the cell length changes are less than 50% of the original length when Rac is uniform highly activated throughout the domain (for reasonable *b* and *δ*).

For the 2D mechanical model, we chose Stokes damping coefficient *γ* = 5.0, viscous damping coefficient *β* = 2.0, Young’s modulus *E* = 1.0, Poisson ratio *ν* = 0.25, Polymerization force density *f_R_* = 0.3/n, switch parameters *s*_0_ = 1 and *s*_1_ = 10, so that changes in the cell area are smaller and slower compared to Rac activity, and so that the cell area changes are around or less than 20% of the original area when Rac is uniform highly activated throughout the domain (for reasonable *b* and *δ*), while maintaining numerical stability.

## Results

### Local perturbation analysis and steady-state behavior of the Rac signaling model

We first use the LPA to approximately map how dynamics of the signaling system in the *bδ*-plane. Results show the *bδ*-plane can be divided into three regions (Figure 2A). (1) A non-polar regime where the only stable solutions to the PDE are homogeneous steady-states which corresponding to a uniform Rac activity level across the cell. (2) A polarizable regime where a sufficiently large spatially heterogeneous stimulus can result in the formation of a polarized pattern. (3) A Turing regime where the homogeneous steady-states are linearly unstable leading to polarization. Numerical simulations of the full PDE system verify the presence of these different regimes. Figure 2B-E illustrate the stable steady-state Rac distributions at the four marked points in Figure 2A. In the case of point C, random noise from the homogeneous steady-state is sufficient to generate patterning (here we used a small amplitude sine function). For D, a large heterogeneous perturbation is required, consistent with the LPA prediction. Neither B nor E is capable of polarizing.

### Proof of concept: fixed domain, 1D model with manually varied tension

The bifurcation structure in Figure 2A suggests a possible mechanism for adaptation in the presence of high levels of signaling. Consider a large chemoattractant that over-stimulates a cell, leading to uniform Rac activation and protrusions across the periphery. In modeling terms, this would correspond to an increase in the basal activation parameter (*b*) as illustrated in Figure 3A. This broad protrusion would lead to an increase in tension (and the parameter *δ*), moving the model cell back into the polarization regime. In this way, uniform high levels of stimulation would initially lead to cell wide protrusion that later gives way to polarization as observed experimentally. In the coming sections, we explore this idea in more detail.

**Figure 3:**
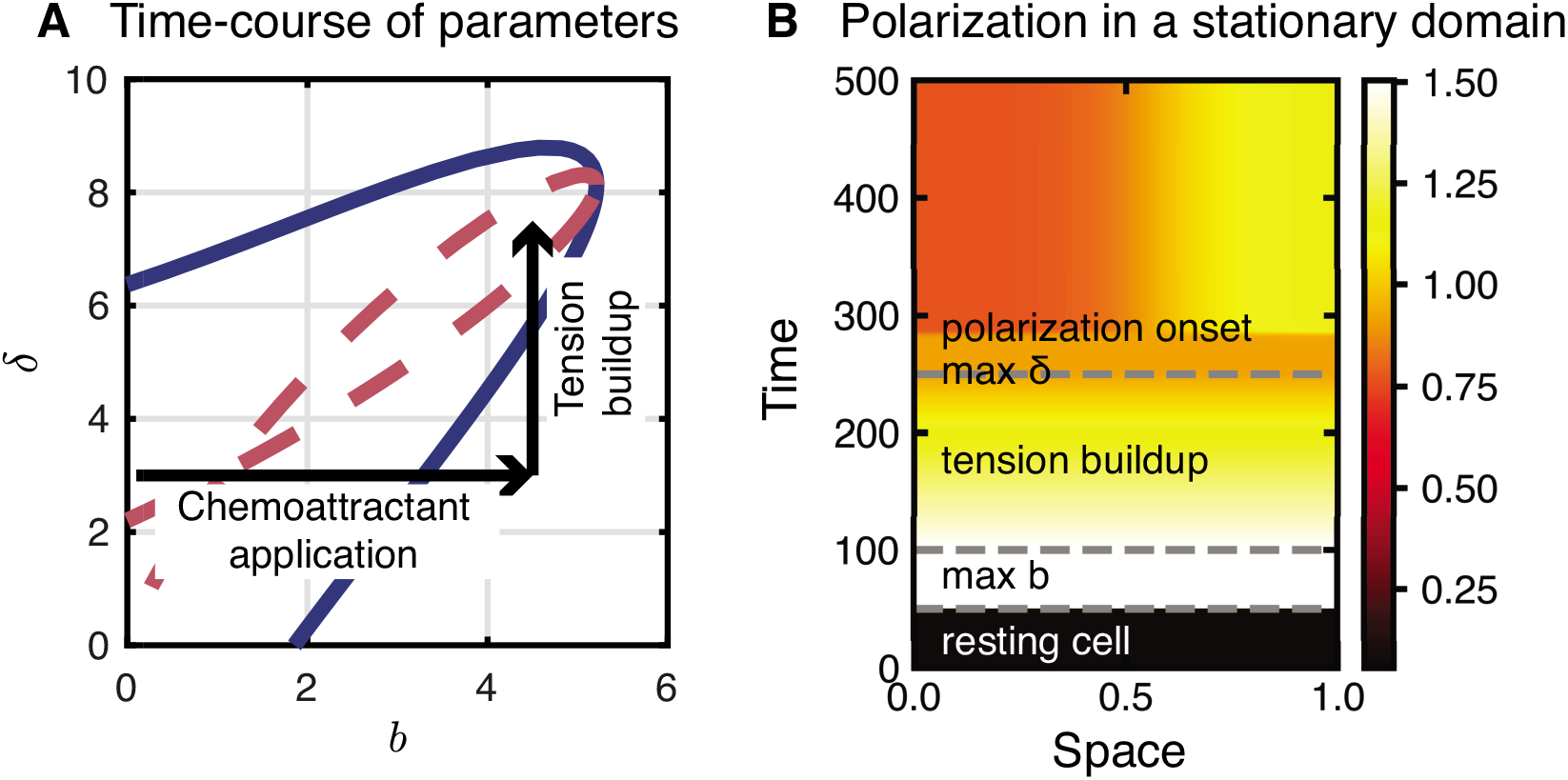
Time-dependent parameters on a stationary domain mimic observed neutrophil behavior. **A** Trajectory of time-dependent parameters in the *bδ*-plane mimicking uniform activation followed by increased tension-mediated inactivation. The blue and dashed red curves are the same as in Figure 2A. Note that for a wide-range of parameter values, a change in *δ* (moving vertically across the bifurcation diagram) can push the cell into the polarizable or Turing regime in response to a change in chemoattractant levels which modulate the activation of Rac through the parameter b (moving horizontally). **B** Kymograph of Rac activity in a stationary domain simulation for these time dependent parameters. The cell is at rest for *t* < 50 with *b* = 0.1, *δ* = 3 (other parameters as in Figure 2). For 50 < *t* < 100 (“max *b*”), the cell is stimulated with uniform chemoattractant *b* = 4.5, and jumps to the high Rac homogeneous steady-state. For 100 < *t* < 250 (“tension buildup”), *δ* increases linearly in time: *δ*(*t*) = 3 + 4.5/150(*t* − 100) to reach a maximum of 7.5 at *t* = 250 (“max *δ*”). As a result of the increasing *δ*, the system re-polarizes with parameters in the Turing regime (“polarization onset”). Polarization results from numerical error in the numerical solution of the PDE system; however, in living cells stochastic effects would drive polarization. See Movie 1.

To begin, we first simulate the 1D spatial version of this model on a fixed domain. Since the domain is fixed, we do not model cell size change and tension changes explicitly but rather temporally change parameters to account for those effects. To mimic uniform stimulation followed by tension build up, we temporally vary the model parameters in two phases. First, *b* is increased to mimic stimulation, as indicated by the horizontal arrow on Figure 3A. Second, we gradually increase *δ* following the vertical arrow. Figure 3B shows the results of this simulation. The initial resting cell becomes highly uniformly stimulated upon onset of stimulation. As tension builds up, uniform Rac activation levels decrease. Finally, without the application of any further perturbation, the model cell polarizes. This suggests the mechanism of expansion followed by tension build up may facilitate adaptation to high signaling levels. This modeling experiment acts as a proof of concept of the mechanosensing-adaptation hypothesis.

### 1D mechanochemical moving-boundary model demonstrates tension mediated adaptation

Using the 1D mechanochemical model, we mimicked the simulation depicted by the arrows in Figure 3A. A cell at rest (uniform Rac activity and stable length) is stimulated by suddenly increasing *b*. From this point on, all cellular changes are driven by the mechanochemical model. Figures 4B and C demonstrate the results of two simulations that yield distinct results. In Figure 4B, the high levels of activation lead to a period of uniform expansion. As predicted by prior simulations, after the cell has sufficiently expanded, tension increases lead to polarization. This confirms the plausibility of this mechanosensing-adaptation hypothesis in a 1D deforming cell model.

**Figure 4:**
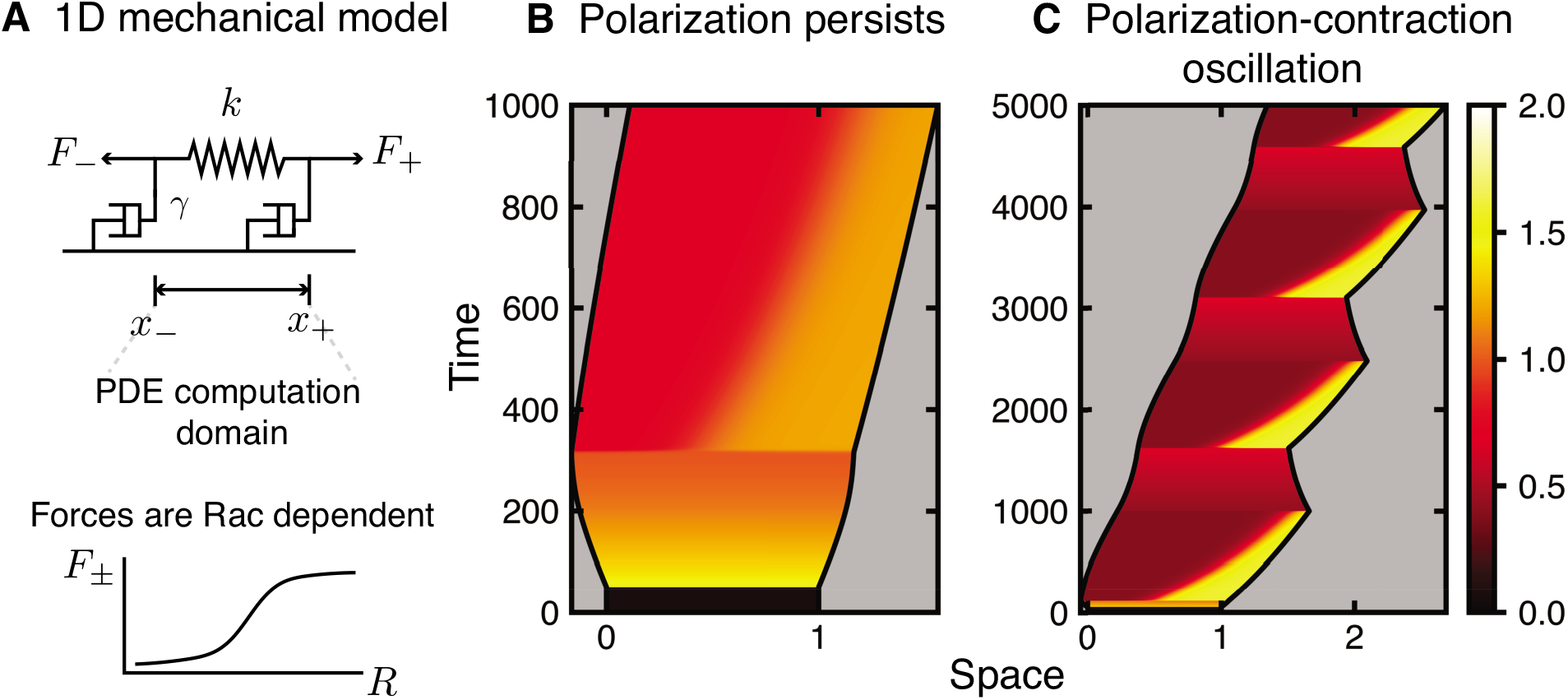
1D mechanochemical model and tension-mediated cell polarization. **A** The cell is modeled as a 1D elastic spring (spring constant *k*) and viscous damping (damping constant *μ*). The PDEs are solved in the domain Ω(*t*) = [*x*_−_(*t*), *x*_+_(*t*)] which varies over time due to forces *F*_±_ that depend on Rac activity at the cell ends (*x*_±_(*t*)). **B**. Uniform stimulation of Rac leads to expansion followed by polarization, consistent with neutrophil observations. Black lines show *x*_−_(*t*) and *x*_−_(*t*) and color shows Rac activity. For this simulation, the cell is initialized at rest at time *t* = 0. The basal activation rate parameter jumps from *b* = 0.1 to *b* = 4 at time *t* ≥ 50. Tension feeds back onto inactivation according to *δ*(*t*) = *δ*_0_ + *δ*_1_*T, T* = *L*(*t*) − ℓ_0_, *δ*_0_ = 3, *δ*_1_ = 1, ℓ_0_ = 1. Other parameters are *f_R_* = 0.001, *s*_1_ = 10, *s*_0_ = 1, *γ* = 1, *k* = 0.001, *c* = 5, *n* = 6, *R_T_* = 2, *D* = 0.01, *D_i_* = 10. Initial conditions are *R*(*x*, 0) = 0.05645 and *R_i_*(*x*, 0) = *R_T_* − 0.05645. See Movie 2. **C** Similar simulation to B but with the basal activation rate increasing from *b* = 0.1 for *t* < 50 to *b* = 1.5 for *t* ≥ 50. Results indicate cyclical phases of polarization and contraction. Other parameters and initial conditions are as in B. See Movie 3.

The second simulation (Figures 4C) illustrates a different type of dynamic that occurs at lower levels of stimulation. Here, the cell undergoes oscillatory phases of polarized protrusion followed by loss of polarity and contraction. In this case, the levels of stimulation are not sufficiently high to maintain polarization when tension-mediated inactivation increases. We note that this polarization-contraction oscillation may be a consequence of the lack of explicit contractile signaling along with the simplified nature of cellular mechanics used in this model. Thus while we include it here for completeness, it is not the focus of this study.

### Bifurcation and parameter space analysis

To supplement our proof of concept example simulations in Figure 3 and 4, we next sought to understand the relationship between cell length changes, chemoattractant levels in the environment (modeled by the parameter *b*), and the resulting cell behaviors: polarized, polarization-contraction oscillation, and non-polarized. First, we performed a LPA analysis of this system where length is itself a parameter. In this approach, length affects the model in two ways: (1) it controls tension and (2) leads concentration-dilution effects resulting from a fixed total amount of GTPase being distributed on a larger cell surface area.

We used LPA to map the cell behavior in the *bL*-plane (Figure 5A). Results demonstrate that the cell can exist in either a non-polar state, stimulus-induced polarized state, or in a linearly unstable regime where polarization results from noise (Turing regime), depending on the current cell length, *L*, and the given basal activation rate *b*. Using this result, we can now map the dynamics of the deforming 1D model in terms of length rather than *δ*.

**Figure 5:**
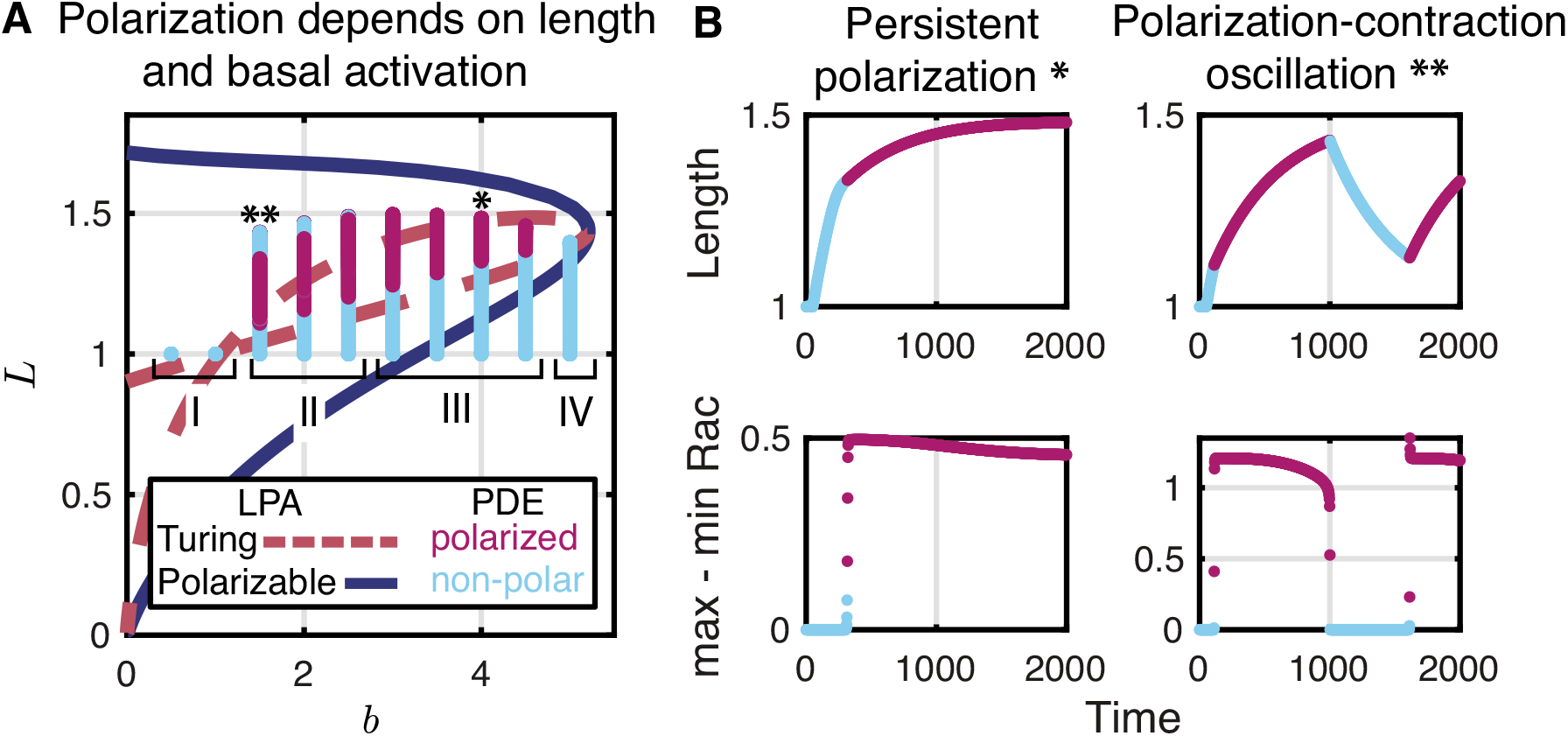
Analysis of length dependence of dynamics. **A** Local perturbation analysis (LPA) of the fixed domain system (*L* is thus a parameter, not a variable) reveals regimes of non-polar, stimulus-induced polarization (inside the blue curve), and Turing instability (i.e., polarization; inside the red dashed loop) in the *bL*-plane. Blue and magenta curves illustrate length *L* trajectories from PDE simulations for various *b* values. As the length changes due to Rac activity (dependent on mechanical parameters), the cell can be in a polarized (magenta) or non-polar (blue) state. Once the cell’s length is in the Turing regime, polarization may occur. Regions I-IV denote the characteristic behavior for a range of *b* values: I Non-polar, resting cells; II Polarization-contraction oscillation; III Persistent polarization; IV Non-polar, expanded cells. **B** Plots of cell length (top row) and polarity strength (bottom row) for two example simulations of the coupled model in regimes III (*b* = 4, left column, labelled * in A) and II (*b* = 1.5, right column, labelled ** in A) respectively. Top row: length as a function of time. Bottom row: difference between maximum and minimum Rac activity as a function of time (a proxy for polarization). Color indicates polar (magenta) or non-polar (blue) Rac activity.

In Figure 5A, we superimpose the length variations of the simulated 1D cell onto the bifurcation diagram in the *bL*-plane. We simulated several 1D deforming cells with different values of *b* (exactly as in Figure 4B and C). Figure 5A illustrates the trajectories of these in length-space as blue and magenta trajectories. Blue indicates that the cell is apolar at that point in the trajectory while magenta indicates polarity. For small *b* values (region I), the Rac activity remains sufficiently low that the cell’s length does not change and the cell remains non-polar. For intermediate *b* values (region II), the Rac activity is sufficient to induce length changes, resulting in Rac polarization. However, there is sufficient Rac activity to increase the cell length beyond the Turing regime, and polarization is lost so that the cell contracts. When the length returns to the Turing regime, spontaneous polarization reoccurs and the cell again lengthens, inducing a polarization-contraction oscillation (as shown in Figure 4C). Quantification of cell length and level of polarity (top and bottom panels respectively of Figure 5B) further illustrate the link between polarized growth and apolar contraction.

For larger intermediate values of *b* (region III), the cell lengthens and polarization persists. In this case, the length increases monotonically before approaching a constant value, with a persistent polarity. Finally, for large *b* values (region IV), stimulation is too strong for polarity to occur. Cells in this region remain non-polar and expanded. We do note that receptor saturation or other factors could limit the maximum level of *b* since Rac activation is downstream of a receptor mediated signaling cascade. Thus if the maximum level of receptor signaling is limited, it is possible that no level of external stimulation would perpetually overstimulate the cell.

### Verification of the mechanosensing-adaptation hypothesis in a 2D continuum mechanics model

Using the 2D mechanochemical model, we repeated the simulation depicted by the arrows in Figure 3A and shown using the 1D model in Figure 4B. In Figure 6, we perform the same simulation experiment where an initially at rest, circular cell is uniformly stimulated with high levels of chemoattractant. Results show that after stimulation, the cell initially expands isotropically to a point and subsequently polarizes and begins migrating. These results confirm that tension-signaling feedback still leads to adaptive polarization when embedded in a 2D geometry.

**Figure 6:**
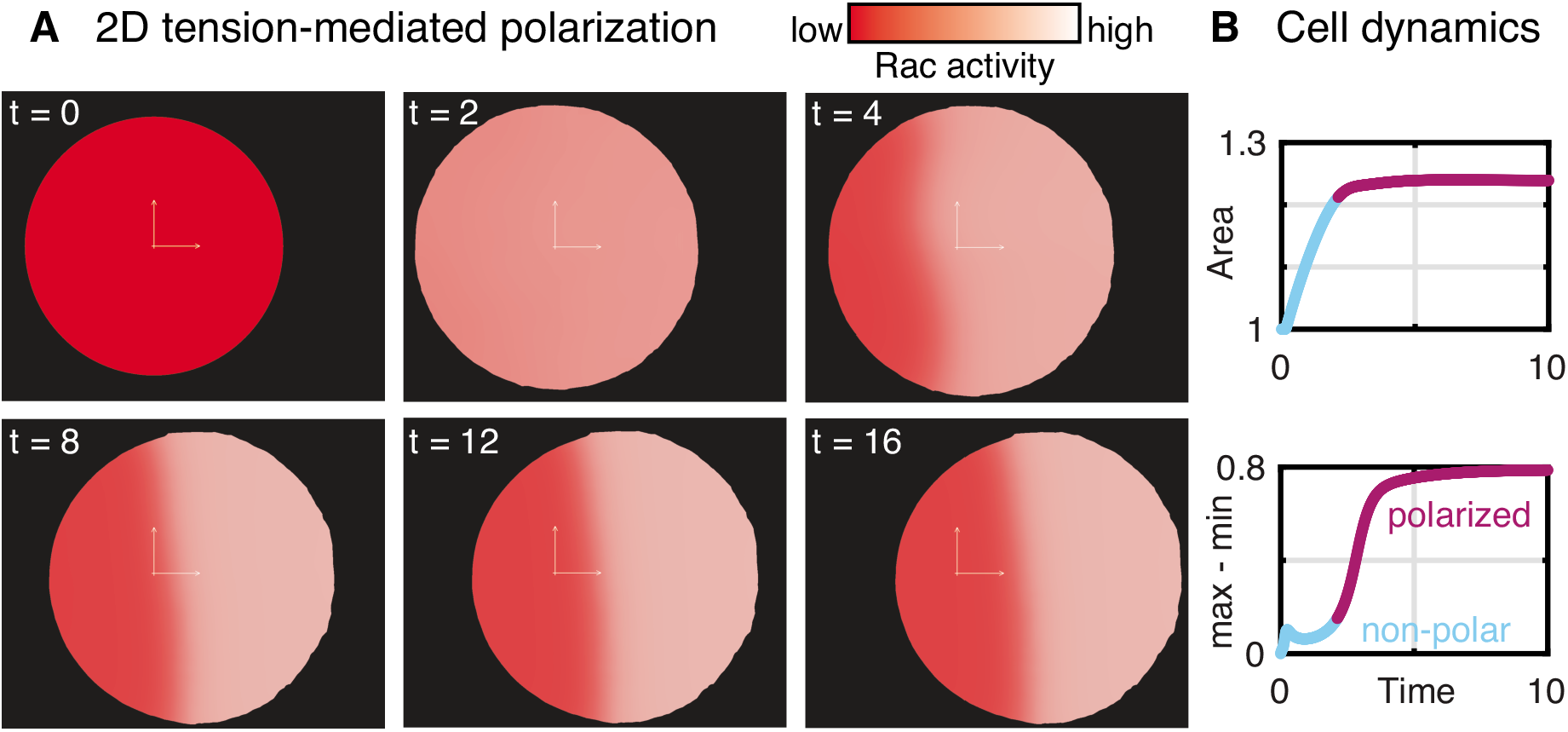
Cell polarization in the 2D mechanochemical model. **A** Snapshots of a 2D simulated cell. The cell is resting when exposed to chemoattractant at *t* = 0, expands due to increased Rac activity, and subsequently polarizes and migrates due to increased tension. Lighter color indicates higher Rac activity. Inset axes show the origin and provide scale; each arm is 0.2 units long. **B** Cellular dynamics. Top row: Area as a function of time. Bottom row: difference between maximum and minimum Rac activity as a function of time (a proxy for polarization). Color indicates polar (magenta) or non-polar (blue) Rac activity. Tension feeds back onto inactivation according to *δ*(*t*) = *δ*_0_ + *δ*_1_*T, T* = *A*(*t*) − *A*_0_, *δ*_0_ = 3, *δ*_1_ = 1, *A*_0_ = 1. We apply a small chemoattractant gradient to ensure polarization along the *x*-axis according to *b*(*x*) = *b*_0_ + *b*_1_*x, b*_0_ = 3, *b*_1_ = 0.1 (the cell is fixed to the *x*-axis at all nodes between *y* = ±0.001 for numerical stability). Other kinetic parameters are *D_x_* = *D_y_* = 0.01, *D_ix_* = *D_iy_* = 100, *c* = 5, *n* = 6, *R_T_* = 2, *f_R_* = 0.3/*n* (*n* is the number of finite element nodes on the boundary), *s*_1_ = 10, *s*_0_ = 1, *h* = 1, *E* = 1, *ν* = 0.25, *γ* = 5, and *β* = 2. Initial conditions are *R*(*x, t*) = 0, *R_i_*(*x*, 0) = 2, *u*(*x*, 0) = 0. No-flux boundary conditions are used for Rac activity, displacement on the *x*-axis does not change in the *y*-direction, and 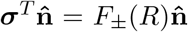 along the boundary. See Movie 4.

## Discussion

In this study, we coupled a reaction-diffusion model of Rac dynamics to a minimal model of cell mechanics to illustrate that the two-way feedback from membrane tension to Rac activity allows cells such as human neutrophils to adapt to saturating levels of chemoattractive environment and maintain cell polarity. We first used the Local Perturbation Analysis (LPA) to map the parameter regimes of behavior of the wave-pinning model with respect to the basal-activation (*b*, which is altered by stimulation) and deactivation rates (*δ*, which is altered by tension). Given the shape of polarizable region in the *bδ*-plane, we hypothesized that an increase in the basal-activation that pushes the cell into an over-activated state can be counter-balanced by a commensurate increase in the deactivation rate *δ* arising from increases in cell tension resulting from protrusion. We then simulated the neutrophil’s response to stimulation from high-levels of uniform chemoattractant behavior using 1D and 2D deformable cell models where signaling influences cell size and tension, and vice versa. Results confirmed that the interaction between tension and signaling can lead to polarization adaptation to high signaling levels. Finally, we coupled the LPA analysis with these full model simulations to demonstrate that the adaptive polarity that arises after uniform stimulation is the result of the cell passing through a bifurcation in the GTPase system. Thus, cell area changes that result from over-stimulation force the cell back into a polarizable state by counter-balancing that stimulation. These results broadly support the hypothesis that mechanosensing can act as an adaptive response to maintain polarity in migrating cells. Further, they are consistent with the conclusions of experimental observations connecting cell mechanics and cell signaling [3–5].

Numerous studies have used differing computational approaches to study the consequences of mechanics, geometry, and tension on cell dynamics such as: [69, 70] modeled competition between separate front and rear cellular compartments, [46, 47, 49, 51, 54, 55] used phase-field simulations incorporating feedback from tension and other physical parameters, [53, 71, 72] used cellular Potts model simulations, [73–75] used level-set methods, [45, 76, 77] used immersed-boundary models, [48, 78] used free-boundary models, and [50, 79] used stochastic methods. Our work differs from these numerous prior studies in that we are studying how the interplay between mechanics and signaling facilitate adaptation. In this sense, our work is most similar to that of Buttenschön et al. [65], who considered the wave-pinning model coupled to a similar 1D mechanical model. Their work did not however incorporate feedback from mechanics to signaling. Zmurchok et al. [80] investigated the consequences of this feedback. However, as this study was primarily concerned with collective, multi-cellular dynamics, each cell was treated as a well-mixed point entity (i.e., polarity was not possible) and adaptation was not studied. Thus, while our mechanochemical modelling fits into this larger body of literature, the approach and intent are distinct. Whereas many of these prior studies investigated the role of mechanics or signaling on cellular behavior, we specifically investigate their interaction. Furthermore, here we investigate the influence of these interactions on adaptation rather than the migratory process itself.

This study does have limitations. We chose to allow cell mechanics to affect the deactivation rate (*δ*). This is likely an over-simplification. It is possible that tension could influence activation in an inverse way, though this would likely lead to similar qualitative results. We also chose to study the simplest model for Rac signaling, and did not explore more complicated variants that include multiple GTPases [26, 31, 33] or additional feedback networks [72]. This was motivated by both simplicity and the fact that the interaction between Rac and tension has been well-studied experimentally [3–5]. We also defined cell tension as a global quantity despite evidence suggesting that membrane tension is more localized than previously thought [81] and that cells such as keratocytes generate tension gradients [82, 83]. In our study, however, we are concerned primarily with how cells respond to uniformly high stimulation that would be expected to, broadly speaking, lead to global increases in tension due to protrusion. Lastly, we used the simplest mechanical models appropriate for cell mechanics (an over-damped elastic spring in 1D and linear viscoelasticity in 2D). While this is sufficient for our purpose, a more detailed model of cell mechanics would be required to study 2D migration itself (as has been done by many others [44–54]).

Despite these limitations, this study demonstrates that mechanosensing can act as an adaptive response to maintain polarity in environments with high levels of chemoattractant. We confirm that the feedback for establishment of polarity via membrane tension can be described using a wave-pinning model of Rac signaling. While we have assumed that membrane tension is a global quantity that feeds back into Rac signaling dynamics, we note that the underlying signaling network is responsible for the adaptive response. Any signal that acts on Rac dynamics in a similar way could induce a similar response (calcium influx, for example, as suggested by Shi et al. [81]).

## Data availability

The code used to produce the Figure 2–5 and Figure 6B is available as an archived GitHub repository at https://doi.org/10.6084/m9.figshare.11916495. Code for the 2D simulations (Figure 6) can be made available upon request.

## Author contributions

CZ and WRH designed the research. CZ and JC performed the research and analyzed data. All authors wrote the paper.

## Acknowledgements

This work was supported by a National Science Foundation grant DMS1562078 (to WRH), a Natural Sciences and Engineering Research Council of Canada (NSERC) Postdoctoral Fellowship Award (to CZ). The authors acknowledge Christopher Bradley for helpful discussions on numerical implementation of the coupled mechanics and reaction-diffusion simulations and Andreas Buttenschön for helpful discussions of the 1D model.

## Supporting Citations

References [84–86] appear in the Supporting Material.

